# The DNA Replication Checkpoint Targets the Kinetochore for Relocation of Collapsed Forks to the Nuclear Periphery

**DOI:** 10.1101/2024.06.17.599319

**Authors:** Tyler Maclay, Jenna Whalen, Matthew Johnson, Catherine H. Freudenreich

**Affiliations:** Department of Biology, Tufts University, Medford, MA 02155

**Keywords:** Nuclear Pore, Collapsed Fork, DNA Replication Checkpoint, Damage Induced Microtubule, DNA Repair, Nuclear Periphery, Rad53, Dun1, TNR, Non-B DNA, Huntington’s Disease, CAG Repeat

## Abstract

Hairpin forming expanded CAG/CTG repeats pose significant challenges to DNA replication which can lead to replication fork collapse. Long CAG/CTG repeat tracts relocate to the nuclear pore complex to maintain their integrity. Forks impeded by DNA structures are known to activate the DNA damage checkpoint, thus we asked whether checkpoint proteins play a role in relocation of collapsed forks to the nuclear periphery in *S. cerevisiae*. We show that relocation of a (CAG/CTG)_130_ tract is dependent on activation of the Mrc1/Rad53 replication checkpoint. Further, checkpoint-mediated phosphorylation of the kinetochore protein Cep3 is required for relocation, implicating detachment of the centromere from the spindle pole body. Activation of this pathway leads to DNA damage-induced microtubule recruitment to the repeat. These data suggest a role for the DNA replication checkpoint in facilitating movement of collapsed replication forks to the nuclear periphery by centromere release and microtubule-directed motion.

**Highlights:**

- The DNA replication checkpoint initiates relocation of a structure-forming CAG repeat tract to the nuclear pore complex (NPC)
- The importance of Mrc1 (hClaspin) implicates fork uncoupling as the initial checkpoint signal
- Phosphorylation of the Cep3 kinetochore protein by Dun1 kinase allows for centromere release, which is critical for collapsed fork repositioning
- Damage-inducible nuclear microtubules (DIMs) colocalize with the repeat locus and are required for relocation to the NPC
- Establishes a new role for the DNA replication and DNA damage checkpoint response to trigger repositioning of collapsed forks within the nucleus.

## Introduction

The nuclear periphery (NP) has emerged as an important site for the maintenance of genome stability when normal replication is impaired. Several types of replication stress have been shown to provoke relocation to the nuclear periphery in both yeast and human cells. These include drugs known to stall or collapse replication forks such as hydroxyurea combined with methyl methanesulfonate (MMS) in yeast ^1^, expanded CAG repeats that form secondary structures ^2^, the polymerase inhibitor aphidicolin in human cells ^3^, protein mediated stalls by replication fork barriers (RFBs) ^4,5^, and replication stalls within telomeres ^6,7^. Some persistent types of DNA damage such as double-strand breaks (DSBs) lacking a readily available template for repair or in sequestered heterochromatin domains and eroded telomeres also relocate the nuclear periphery (for review, see ^8–10^).

We previously elucidated a detailed pathway for how collapsed forks caused by an expanded CAG/CTG tract of 130 repeat units (abbreviated (CAG)_130_) relocates to the *S. cerevisiae* nuclear pore complex (NPC) in late S phase in a manner dependent on replication ^2,11^. CAG/CTG repeats over 35 repeat units have a propensity to form hairpin structures that can impair replication leading to fork collapse ^12,13^. Defects in relocation to the NPC result in increased chromosome end loss and repeat instability, implicating this pathway in protecting against fork breakage and inaccurate repair ^2^. The DNA structure induced relocation pathway has many similarities with pathways that have been described for persistent or heterochromatic DSBs and eroded telomeres, but also several differences (reviewed in ^8^). We determined that relocation of collapsed forks caused by CAG repeats was dependent on resection, the binding and SUMOylation of key fork associated repair proteins (RPA, Rad52, Rad59, Smc5) by the Smc5-6 associated SUMO ligase Mms21, and interaction with the SUMO interacting motifs (SIMs) of Slx5 which targets the collapsed fork to the NPC ^11^.

In addition to the SUMOylation requirements, we wanted to investigate other processes that could be involved in relocation of collapsed forks caused by CAG repeats to the NPC. One important pathway known to respond to stalled and collapsed forks is the DNA damage response. The DNA damage response can be subdivided into two pathways: the DNA replication checkpoint (DRC) and the DNA damage checkpoint (DDC), defined by their mediator proteins Mrc1 or Rad9, respectively. Our previous findings showed that expanded CAG repeats activate the DDC ^14^, and long range resection is required for (CAG)_130_ relocation ^11^, which would produce ssDNA to stimulate DDC activation. In addition, members of the DDC and the DRC were important for maintaining repeat stability and preventing chromosome breaks at long CAG tracts ^15–17^.

The checkpoint is composed of several sensor and mediator components that activate downstream effectors, reviewed in ^18,19^ and summarized in (Figure 1A). Sensor proteins identify processed DNA lesions, such as resected DNA substrates, and then recruit other proteins to the damage site. In *S. cerevisiae*, the sensor Ddc2 (hATRIP) interacts with RPA bound to ssDNA to recruit the kinase Mec1 (ATR). Mec1 is also activated by the PCNA-like 9-1-1 clamp (Ddc1, Rad17, Mec3 in budding yeast) which is loaded by the Rad24-RFC complex onto ds/ssDNA junctions. Downstream effector proteins such as Rad53 (hChk2) and Chk1 elicit the DNA damage response through further phosphorylation signaling. At the replication fork there is an additional Rad53 activation mechanism involving the mediator, Mrc1 (hClaspin), which travels with the replisome and is phosphorylated upon replisome uncoupling ^20,21^. Another pathway more specific to DSBs involves Tel1 (hATM), which senses DNA ends through its interaction with the yeast MRX (hMRN) complex. Finally, the yeast mediator protein Rad9 (h53BP1) amplifies Rad53 phosphorylation. Rad9 is recruited to histone modifications including methylated H3 (H3-K79) and damage-specific H2A-S129 phosphorylation, which can occur at either collapsed forks or DSBs. Due to the different DNA damage sensors, identification of the checkpoint pathway that is activated can give insight into the lesion or lesions that are created by a particular type of DNA damage.

**Figure 1:**
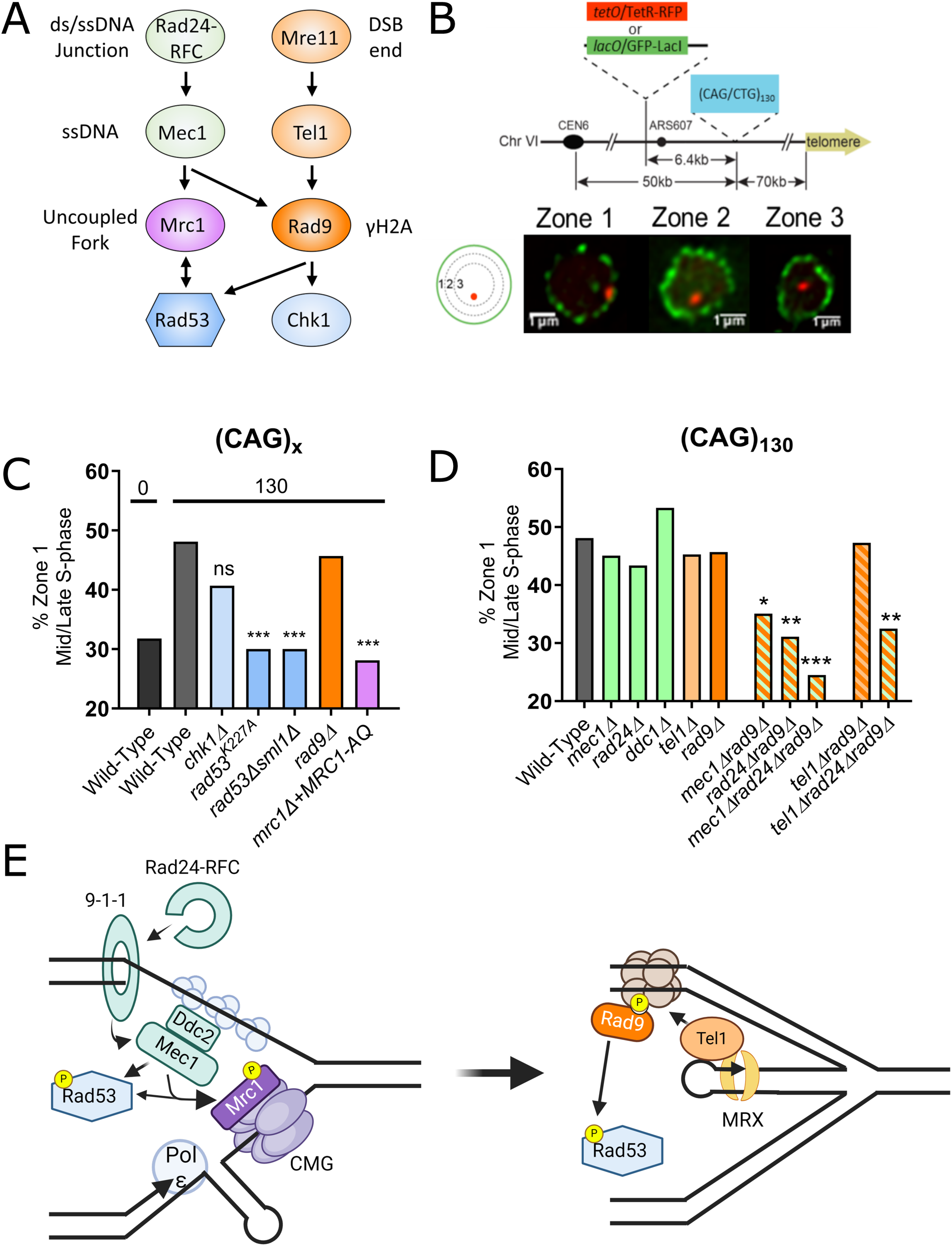
Relocation of (CAG)_130_ locus to the nuclear periphery depends on the activation of the DNA replication or DNA damage checkpoint. A) Flow chart of the DNA damage checkpoint. B) Yeast chromosome VI containing integrated 130 CAG/CTG repeat tract ((CAG)_130_) and a *lacO* or *tetO* array that binds the LacI-GFP or TetR-RFP protein, respectively. Mid/late S phase cells were imaged, and the location of the foci were scored into one of three zones of equal area, using the GFP-Nup49 signal to mark the nuclear periphery. Representative images are shown. C) Percent of cells with LacO loci in zone 1 for mid/late S-phase cells. Cells have either (CAG)_0_ or (CAG)_130_ as marked. D) Percent of zone 1 foci for mid/late S-phase cells for indicated strains. All strains have (CAG)_130_. E) Model illustrating activation of the DRC in response to an uncoupled fork (left) or activation of the DDC following fork reversal and recruitment of the MRX complex (right). (*) P≤0.05, (**) P≤0.01 (***) P≤0.001 compared with wild-type (CAG)_130_ strain by Fisher’s exact test. More than 100 cells were analyzed per strain. See Table S1 for the exact number of cells analyzed, percentages, and P-values.

The relocation of persistent DSBs in *S. cerevisiae* and heterochromatic DSBs in *Drosophila* required both Mec1/Tel1 and ATR/ATM, respectively ^1,22,23^. In human U2OS and IMR90 cells that were treated with aphidicolin to inhibit polymerases, ATR inhibition prevented the localization of stalled replication forks to the nuclear periphery ^3^. In contrast, single deletion of either Mec1 or Tel1 did not prevent the relocation of an expanded (CAG)_130_ repeat to the NPC ^2^, leaving it an open question whether the checkpoint is involved in the relocation of DNA damage caused by DNA structures to the nuclear periphery, and if so what pathway is involved. In yeast, Mec1/ATR is not only required for DSB relocation, but has been linked to increasing chromatin mobility in response to DSBs ^24^. Yet an expanded (CAG)_130_ repeat not only failed to increase mobility of a chromosome harboring this fork-stalling element, it decreased mobility due to the added constraint of NPC interaction ^2^. Therefore, it was not clear whether the checkpoint pathway would be involved in relocation of DNA structure-mediated collapsed forks to the NPC, and if so, in what manner.

In this study, we show that the activation of the effector kinase Rad53 by phosphorylation of the replisome associated Mrc1 protein is needed for the relocation of a (CAG)_130_ tract to the nuclear periphery. We identified that the Rad53 target Dun1 is also required for relocation. Through investigating several Dun1 substrates, we determined that the phosphorylation of the kinetochore protein Cep3 at serine 575 is the important target to allow for repositioning of the DNA structure-induced collapsed fork in the nucleus. We investigated several possible mechanisms by which Cep3 phosphorylation could act and identified the disruption of centromere attachment to the spindle pole body (the yeast microtubule organizing center which is functionally equivalent to the mammalian centrosome) as a critical factor. In contrast, Cep3 phosphorylation-dependent increases in global chromosome movement were not detectable. Finally, we show that damage-induced microtubules (DIMs) ^25^ are formed in response to damage at the CAG tract, colocalize with the CAG tract locus, and are dependent on this checkpoint pathway. We speculate that the phosphorylation of Cep3-S575 allows for relocation of collapsed forks to the NPC by relieving centromere constraint and supporting DIM-directed motion towards the nuclear periphery.

## Results

### Relocation of a (CAG)_130_ tract to the nuclear pore complex depends on the activation of Rad53 and Mrc1

We used a previously described zoning assay ^2^ to visualize the CAG repeat in relation to the nuclear periphery. Briefly, 130 CAG/CTG repeats are integrated on chromosome VI 6.4 kb away from a LacO array which can be visualized by binding of GFP-LacI; in some experiments the repeat was placed 6.4 kb from a TetO/mCherry-TetR array (Figure 1B). GFP-Nup49 was used to mark NPCs to allow for visualization of the nuclear periphery. The nucleus was divided into three equal areas to determine the frequency of CAG repeat occupancy in each zone (see Methods). Zone 1 represents the nuclear periphery (where the NPCs are located), zone 3 is the middle of the nucleus, and zone 2 is the area in between (Figure 1B). The presence of the (CAG)_130_ tract caused an increase in the occupancy of the nearby tagged locus in zone 1, from 31% in the (CAG)_0_ control to 48% for the chromosome containing (CAG)_130_ (Fig. 1C). The distribution between zones 1, 2 and 3 was consistent between both wild-type (CAG)_130_ strains, whether the LacO/GFP-LacI or TetO/mCherry-TetR array was used to follow the CAG repeat locus (Figure S1).

Previously, it was shown that single deletions in Mec1 or Tel1 sensor kinases (ATR and ATM in mammals, respectively) do not affect relocation of the (CAG)_130_ tract to the nuclear periphery during S phase ^2^. Simultaneous disruption of Mec1 and Tel1 could not be tested by the zoning assay due to severely abnormal nuclear morphology. As there are many factors involved in the activation of the DNA damage response that can act redundantly (Figure 1A), we investigated if there was any role for the checkpoint in CAG tract relocation by testing the effector kinases. The two effector kinases of the yeast DNA damage checkpoint are Chk1 and Rad53 (reviewed in ^18^). Deletion of Chk1 showed no significant decrease in the frequency the CAG tract was found at the nuclear periphery (Figure 1C). However, deletion of Rad53 (viable with the added deletion of Sml1 ^26^) did result in a significant decrease in (CAG)_130_ occupancy in zone 1. This result supports that the DNA damage response is required for relocation. This was further confirmed by testing *rad53^K227A^*, a kinase-defective mutant of Rad53 that is checkpoint-deficient ^27^. *rad53^K227A^* showed the same significant decrease in relocation as *rad53Δsml1Δ* (Figure 1C). We conclude that activation of the DNA damage response and the kinase activity of Rad53 is needed for relocation of the collapsed fork to the NPC.

There are multiple sensor and mediator proteins that can activate Rad53 in response to different types of DNA damage. We had previously shown that long CAG tracts were especially prone to breakage and exhibited increased fork stalling in the absence of the Mrc1 protein ^17^, which couples polymerase epsilon to the CMG helicase. Mrc1 contains multiple SQ/TQ motifs that are targeted by kinases of the PIKK family, such as Mec1 (hATR), Tel1 (hATM) or Rad53 (hChk2) for phosphorylation upon fork uncoupling ^28,29^. The Mrc1^AQ^ mutant, with its 17 SQ/TQ sites mutated to alanine ^29^, had a modest effect on chromosome fragility but exhibited increased repeat instability ^17^, suggesting a possible role for Mrc1 in sensing forks stalled at the (CAG)_130_ tract. The deletion of Mrc1 had severely abnormal nuclear morphology precluding zoning analysis. However, complementation with the checkpoint-deficient Mrc1^AQ^ mutant abrogated these abnormal morphologies and showed a significant reduction in relocation to the nuclear periphery to a similar level as a Rad53 mutant (Figure 1C). Thus, the checkpoint signal relevant for repositioning to the NPC is channeled through Mrc1, which suggests that the signal originates from an uncoupled replication fork.

### Mec1 and Rad9 act redundantly to sense damage at the (CAG)_130_ tract and signal repositioning within the nucleus

Though single mutations in either of the main checkpoint sensor kinases, Mec1 and Tel1, did not result in a decrease in relocation to the nuclear periphery (see Figure 1D) the presence of the Mre11 protein, which senses and processes DSB ends, is required ^11^. Deletion of the Rad9 mediator protein (h53BP1 ortholog), which acts downstream of Tel1 to facilitate Rad53 autophosphorylation ^30,31^, showed no effect on the relocation of the repeat to the nuclear periphery (Figure 1D). Similarly, deletion of the 9-1-1 clamp component Ddc1 (9-1-1 activates Mec1) or the clamp loader Rad24 also had no effect on the percent of the CAG locus in zone 1 (Figure 1D). These results were somewhat surprising given the important role of Mrc1, which is phosphorylated by Mec1, and our previous findings that both Mec1 and Rad9 were important for preventing breaks and instability at expanded CAG tracts ^15^. Since there is a well-described redundancy between these sensing pathways (reviewed in ^18^), we explored whether there was redundancy in the sensing of lesions to signal relocation of the (CAG)_130_ to the nuclear periphery. Deletion of both Mec1 and Rad9 resulted in a decrease in relocation of the (CAG)_130_ to the nuclear periphery (Figure 1D). In addition, co-disruption of Rad9 and Rad24 also led to a decrease in relocation to the NP, and deletion of all 3 genes (*mec1Δrad24Δrad9Δ*) led to the lowest % zone 1 occupancy (Figure 1D). These effects were not made worse by additional disruption of Tel1 (Figure 1D), suggesting that the role of Tel1 is not independent of Rad9 in this situation. We previously showed that both H2A-S129 and H2A-T126 phosphorylation could be detected at a (CAG)_155_ tract 40 minutes into S phase ^32,33^, which is shortly before relocation occurs ^11^ and could serve to recruit Rad9. Importantly, DNA damage dependent phosphorylation of H2A requires either Mec1 or Tel1 ^34,35^. So, in the absence of Mec1, the recruitment of Rad9 would be expected to be downstream of Tel1 activity. Thus, our data indicate that there are multiple ways to activate Rad53 to enable relocation of the (CAG)_130_ tract: a Tel1/Rad9 mediated pathway, or a Rad24/Mec1 pathway, acting redundantly with the Rad9 pathway. In addition, Mrc1 phosphorylation, which facilitates recruitment of Rad53 ^36^, is an important mediator of Rad53 activation at the CAG-stalled fork.

Altogether, this analysis suggests that there may be multiple types of damage at the CAG tract that can provoke the checkpoint response and are important for nuclear repositioning. Based on our data, repeat mediated checkpoint activation and relocation to the NPC could be due to an uncoupled fork (sensed by Mrc1), a region of single-stranded DNA (sensed by the Ddc2-Mec1 and 9-1-1 complexes), or a DSB end (sensed by the MRX-Tel1-Rad9 pathway). Single-strand DNA could occur at an uncoupled fork, at a reversed fork, or at a post-replicative gap, and a DSB end could result either from fork reversal or fork breakage/cleavage. All of these signals would be present at an uncoupled fork that transitions to either a reversed or broken fork, which we refer to collectively as “collapsed forks” (Figure 1E). Alternatively, there could be a mixture of possible species that occur in different cells.

### Phosphorylation of Cep3 Serine 575 by Dun1 is needed for relocation of collapsed forks to the nuclear pore complex

After determining that activation of the transducing kinase Rad53 by the DNA replication checkpoint is required for (CAG)_130_ relocation, we investigated what targets of Rad53 are needed. Dun1 is a well characterized target of Rad53 (Figure 2A) ^37–40^. A *dun1Δ* showed a significant decrease in CAG tract relocation to the N.P. similar to *rad53Δsml1Δ* and *rad53^K227A^* mutants (Figure 2B). Therefore, Rad53 activation of Dun1 is a crucial step in the pathway leading to repositioning of collapsed forks to the NPC.

**Figure 2:**
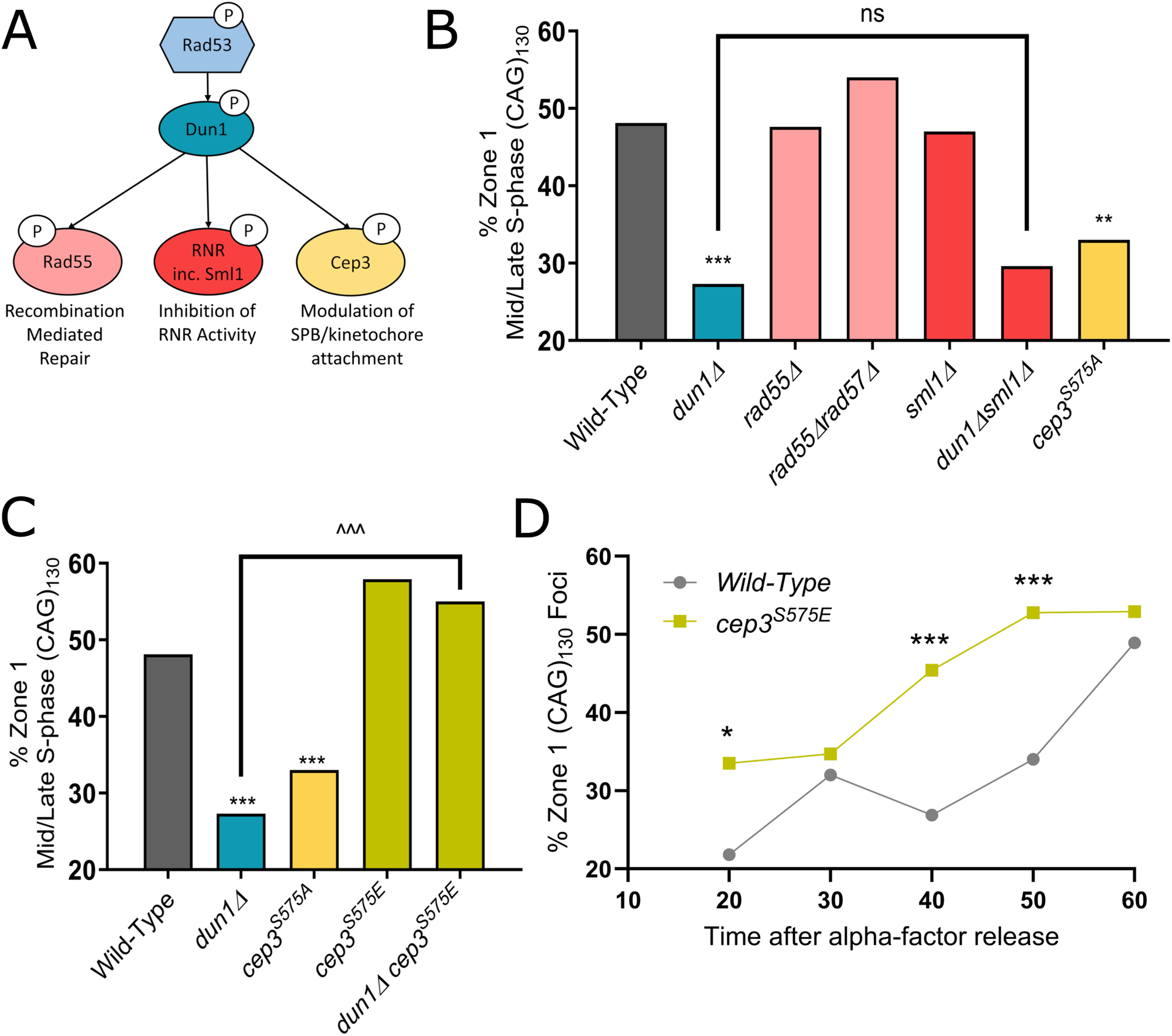
Phosphorylation of Cep3^S575^ by Dun1 is needed for relocation of (CAG)_130_ to the nuclear periphery. A) Diagram of Dun1 targets. The colors used for mutants in this figure correspond with colors used in this diagram. B) Percent of zone 1 foci for (CAG)_130_ mid/late S-phase cells for strains deleted for Dun1 or its targets. C) Percent of zone 1 foci for (CAG)_130_ mid/late S-phase cells for a phosphomimetic mutant of Cep3 (*cep3^S575E^*) individually and in the *dun1Δ* mutant. D) Percentage of zone 1 foci for (CAG)_130_ cells at indicated time points after release from alpha factor arrest in wild-type and *cep3^S575E^*strains. See Table S2 for the exact number of cells analyzed, percentages, and P-values. (*) P≤0.05, (**) P≤0.01 (***) P≤0.001 compared with wild-type by Fisher’s exact test. (^^^) P ≤ 0.001 compared as indicated by Fisher’s Exact Test. More than 100 cells were analyzed per strain. See Table S1 for the exact number of cells analyzed, percentages, and P-values.

Next, we investigated what targets of Dun1 are required for relocation (Figure 2A). Dun1 is required for the phosphorylation of the Rad51 associated repair protein Rad55 ^37^. Phosphorylation of Rad55 allows for it to dimerize with Rad57 and effectively stabilize Rad51 binding to promote more successful strand invasion ^41^. Therefore, Dun1 plays a direct role in recombination mediated repair. However, neither *rad55Δ* nor *rad55Δrad57Δ* mutants showed a decrease in the percentage of CAG tract loci detect in zone 1 (Figure 2B). Dun1 has also been well studied as an activator of ribonucleotide reductase (RNR) activity and functions to regulate nucleotide metabolism in response to replicative stress. Dun1 inhibits Rfx1, Sml1, and Dif1 through phosphorylation which in turn prevents them from downregulating nucleotide pools ^42–44^. Since Dun1 inhibits Sml1 activity, in order to determine if Dun1’s role in relocation is through maintaining nucleotide pools, Sml1 was deleted in the *dun1Δ* strain. If the deletion of Sml1 rescues the defect in CAG tract repositioning in the *dun1Δ* strain, it would indicate that this activity of Dun1 is required. However, *dun1Δsml1Δ* showed a similar percent S phase zone 1 occupancy as *dun1Δ* strains (Figure 2B). We conclude that the role for Dun1 in relocation to the NP is neither to promote recombination mediated repair nor to increase nucleotide pools.

Another phosphorylation target of Dun1 is the kinetochore protein Cep3 ^45^. Cep3 is a DNA-binding protein that recognizes the centromere sequence as a member of the CBF3 kinetochore complex ^46,47^. Phosphorylation of Cep3 at serine 575 (S575) occurs in response to an induced DSB or zeocin-induced DNA damage in a manner dependent on both Rad53 and Dun1 ^45^. The yeast kinetochore interacts with the spindle pole body (SPB), the yeast microtubule organizing center which is functionally equivalent to the mammalian centrosome. Cep3-S575 phosphorylation has been reported to allow for an increase in the dynamics of the SPB-CEN attachment after DSB induction on a different chromosome ^45^. Mutation of serine 575 to a non-phosphorylatable alanine residue resulted in a significant decrease in the CAG locus occupancy in zone 1, showing that Cep3 phosphorylation is required for relocation of the CAG tract to the NPC (Figure 2B).

To confirm that Cep3 phosphorylation at S575 is required for relocation we constructed a phosphomimetic mutant of Cep3 by mutating serine 575 to glutamic acid. The *cep3^S575E^* mutation bypassed the need for Dun1 as the *dun1Δcep3^S575E^* mutant showed a normal level of CAG locus S phase occupancy in zone 1 (Figure 2C). We conclude that phosphorylation of Cep3-S575 mediated by Dun1 is needed for relocation of collapsed forks to the NPC.

To further address the role of Cep3 phosphorylation we assessed the timing of CAG locus relocation to the NP in *cep3^S575E^* cells over the course of an S-phase. As previously reported, in wild-type cells the (CAG)_130_ tract is rarely at the NPC in early S-phase and is maximal in late S phase at 60 min post release from G1 (Figure 2D) ^11^. However Rad52 is found at the repeat locus 40 minutes into S phase ^2^, suggesting that fork collapse has occurred by then and that there is a delay between the occurrence of damage and anchoring at the pore. The *cep3^S575E^* mutation led to a significant increase in zone 1 occupancy of the (CAG)_130_ repeat tract at the 40- and 50-minute time points. (Figure 2D). This suggests that the *cep3^S575E^*phosphomimetic is bypassing the normal signaling pathway, priming the fork for repositioning as soon as enough RPA-coated ssDNA has occurred to recruit Rad52.

### Phosphorylation of Cep3-S575 allows for relocation of collapsed forks to the NPC by modulating kinetochore-SPB attachment, but not by increasing global chromosome movement or activating the spindle assembly checkpoint

It has been shown that forcing transcription through the centromere with a galactose-inducible promoter causes detachment of the centromere from the SPB ^48,49^. Cep3-S575 phosphorylation is needed for kinetochore declustering and increased SPB-CEN dynamics in response to DSBs and has therefore been suggested to relieve attachment of centromeres to the SPB ^45^, though the *cep3^S575A^*mutation was not observed to increase SPB-CEN distance or position with respect to the nuclear periphery after DSB induction ^45,50^. We reasoned that if centromere attachment to the SPB was preventing relocation to the NPC in the *cep3^S575A^* mutant, then forcing transcription through the centromere and thus interrupting that attachment could allow for relocation of the repeat tract in the *cep3^S575A^* mutant. We integrated the *GAL1* promoter upstream of the centromere on chromosome 6 (*pGAL1-CEN6*) ^51^, which is the chromosome containing the (CAG)_130_ tract. Upon galactose induction and transcription through the centromere, the reduced zone 1 occupancy of the CAG locus in the *cep3^S575A^* strain was rescued to normal wild-type levels (Figure 3A). We conclude that phosphorylation of Cep3-S575 may allow for relocation of collapsed forks to the NPC by relieving centromere attachment to the SPB.

**Figure 3:**
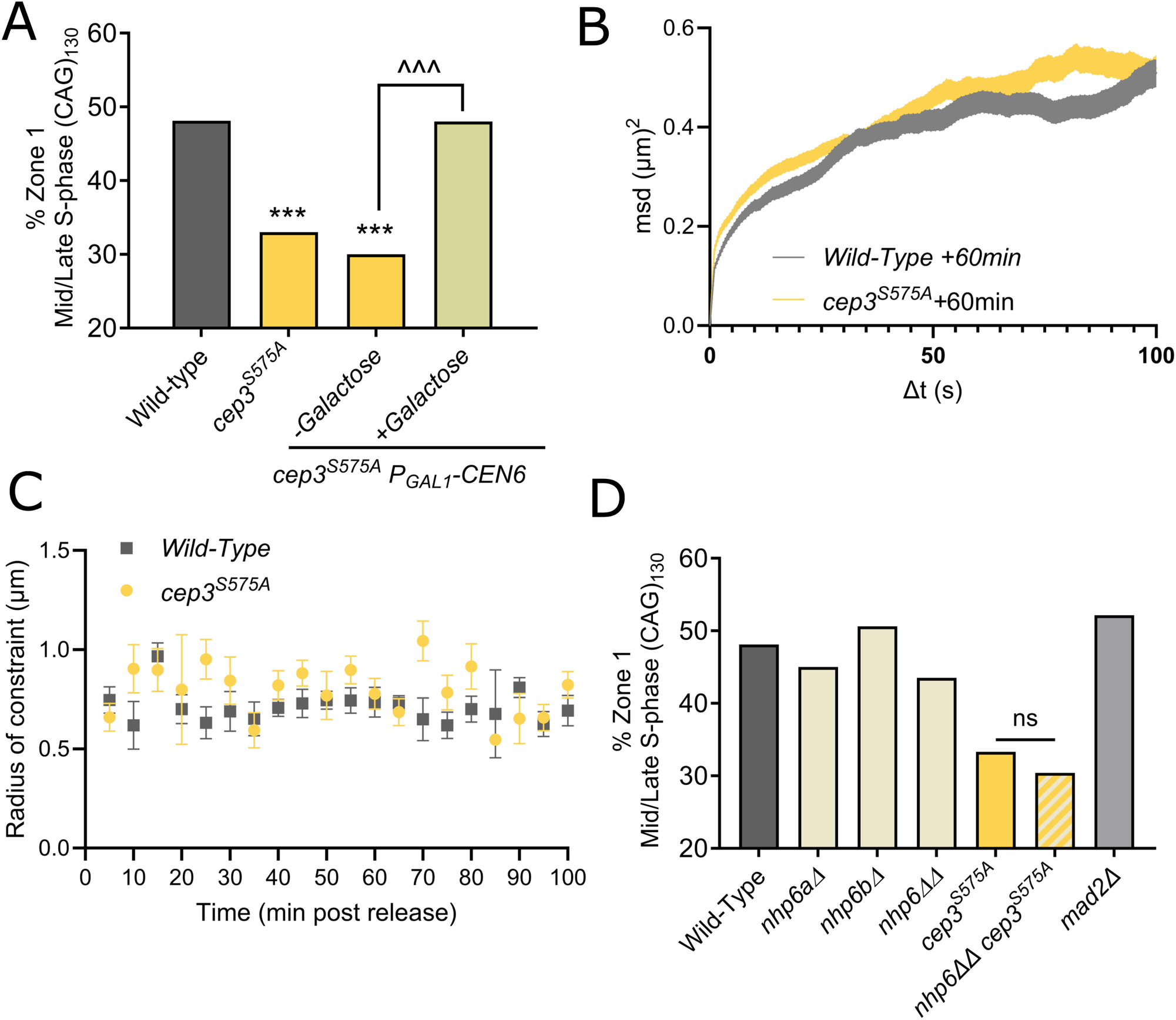
Cep3 phosphorylation allows for (CAG)_130_ relocation through release from centromeric constraint. A) Percent zone 1 foci for (CAG)_130_ mid/late S-phase cells for *cep3^S575A^* and *P_GAL1_-CEN6 cep3^S575A^* strains. Cultures were grown in YP-lactate media then split into media containing either glucose or galactose to induce transcription through CEN6. B) MSD track for Cep3^S575A^ and WT strains containing (CAG)_130_. Cells were imaged every 1.5 seconds for 5 minutes, first Δt =[0,100] seconds shown. Data was captured 60 minutes post-release from alpha factor arrest. The lines represent the 95% Confidence intervals of the mean square displacement at a particular timepoint. C) Summary of radii of constraint calculated for WT and cep3^S575A^ over 100 minutes post release from alpha factor arrest. Points represent calculated Radius of constraint for a 5-minute interval imaged every 1.5s as in B, error bars are standard error of the calculated radius of constraint., specific details in Table S3. D) Percent of zone 1 foci for (CAG)_130_ mid/late S-phase cells >100 cells per strain analyzed, see Table S1 for exact values(***) P≤0.001 compared with wild-type by Fisher’s exact test. (^^^) P= 0.0004 by Fisher’s exact test.

Detachment of the centromere from the SPB has a number of consequences. It has been shown that inactivating a centromere can increase the mobility of that chromosome ^45^. It is also known that induced DSBs increase both local ^24^ and global ^52^ chromosome mobility (reviewed in ^53^). This DSB-induced mobility was shown to be dependent on Cep3 phosphorylation in one study ^45^, but not another ^50^. We have previously published that the (CAG)_130_ tract, in contrast to a DSB, does not increase either local or global chromosome mobility, and instead causes decreased locus mobility relative to chromosomes lacking the repeat tract ^2^. However, those data were collected from unsynchronized cells in S phase, and the locus likely becomes constrained only after anchoring to the NPC. We reasoned that an increase in chromosome mobility due to a collapsed fork at the (CAG)_130_ tract and subsequent Cep3-S575 phosphorylation could be transient. The anchoring at the NPC occurs in a distinct time window in late S phase, between about 55-75 minutes after release from G1 and peaking at ∼60 minutes (Figure 2D and ^11^), therefore any chromosome VI movement could be confined to a short time period preceding NPC anchoring. The radius of constraint, a measurement of locus mobility ^54^, was measured for the (CAG)_130_ locus in cells that were synchronously released from G1 phase over a 100 minute time course segmented into 5 minute intervals. During each 5 minute interval, images were acquired every 1.5 seconds, closely mirroring the conditions used in ^45^, and MSD curves for each 5 minute window were plotted to obtain a radius of constraint. An example of the MSD curve for the first 100 seconds of the 60-minute timepoint is depicted (Figure 3B). For each timepoint, the radius of constraint was calculated based on the plateau reached within Δt = 100s (Figure 3C; Table S3). No difference in mobility of chromosome VI adjacent to the (CAG)_130_ tract was observed between wild-type and *cep3^S575A^* cells (Figure 3C). Though a fast directed motion may not have been detected by this method, this result argues against a local increase in chromosome movement triggered by Cep3 phosphorylation in response to a collapsed fork.

Another driver of DSB mobility is local nucleosome depletion ^50^. Loss of Nhp6, encoded in two loci Nhp6a and Nhp6b, has been shown to reduce histone occupancy on DNA ^55^, and this leads to increased chromosome mobility ^50,56^.To test the mobility hypothesis in a different way, we reasoned that if increasing chromosome mobility by phosphorylating Cep3^S575^ was a requirement for (CAG)_130_ relocation, then causing global nucleosome depletion by deletion of Nhp6a and Nhp6b could rescue the *cep3^S575A^* relocation defect. Deletion of Nhp6a and Nhp6b (*nhp6*ΔΔ) did not cause a decrease in relocation, and when combined with *cep3^S575A^*did not rescue the relocation defect of those cells (Figure 3D). Altogether, we conclude that an increase in chromosome mobility does not explain the importance of Cep3 phosphorylation for relocation of the (CAG)_130_ tract to the NP.

Centromeres unattached to the spindle can activate the spindle assembly checkpoint (SAC) via Mad2 ^57^, and phosphorylation of Cep3 has been shown to activate the SAC ^45^. Therefore, we tested whether activation of the SAC by Mad2 is required for relocation, however there was no relocation defect in *mad2Δ* cells (Figure 3D). We conclude that the role of Cep3 phosphorylation in collapsed fork repositioning to the NP is not due to a defect in SAC activation.

### Expanded CAG repeat loci associate with DIMs in a Dun1 and Cep3-S575 phosphorylation-dependent manner

In yeast, microtubule structures termed “Damage-Inducible Microtubules” or “DIMs” form in response to double stranded breaks in a Rad9 checkpoint-dependent manner ^25^. Interestingly, inactivating a centromere is sufficient to drive formation of DNA damage-inducible microtubules (DIMs) ^25^. Since our data supported that centromere inactivation and Cep3 phosphorylation were important to allow relocation of the CAG locus to the NP, but not through increasing chromosome mobility, we wondered whether DIM formation might be involved. We reasoned that if DIMs were important for relocation to the periphery, then the formation of microtubules in general would be important. To test this, cells were treated for 30 minutes with either 15 μg/mL of the tubulin depolymerizing drug nocodazole or an equivalent amount of the vehicle (DMSO) and the zoning assay was performed. Addition of nocodazole significantly reduced the percentage of the (CAG)_130_ tract localized to the nuclear periphery in mid-to-late S phase cells (Figure 4A). It is known that the minus end-directed microtubule motor Kar3 promotes, though is not strictly required, for DIM formation ^25^. Therefore, we tested whether deletion of Kar3 had an effect on the frequency of (CAG)_130_ locus association with the NP. In *kar3Δ* cells in S phase, relocation of the repeat to zone 1 was reduced (though not eliminated; Figure 4B), in line with the reported *kar3Δ* effect on DIMs.

**Figure 4:**
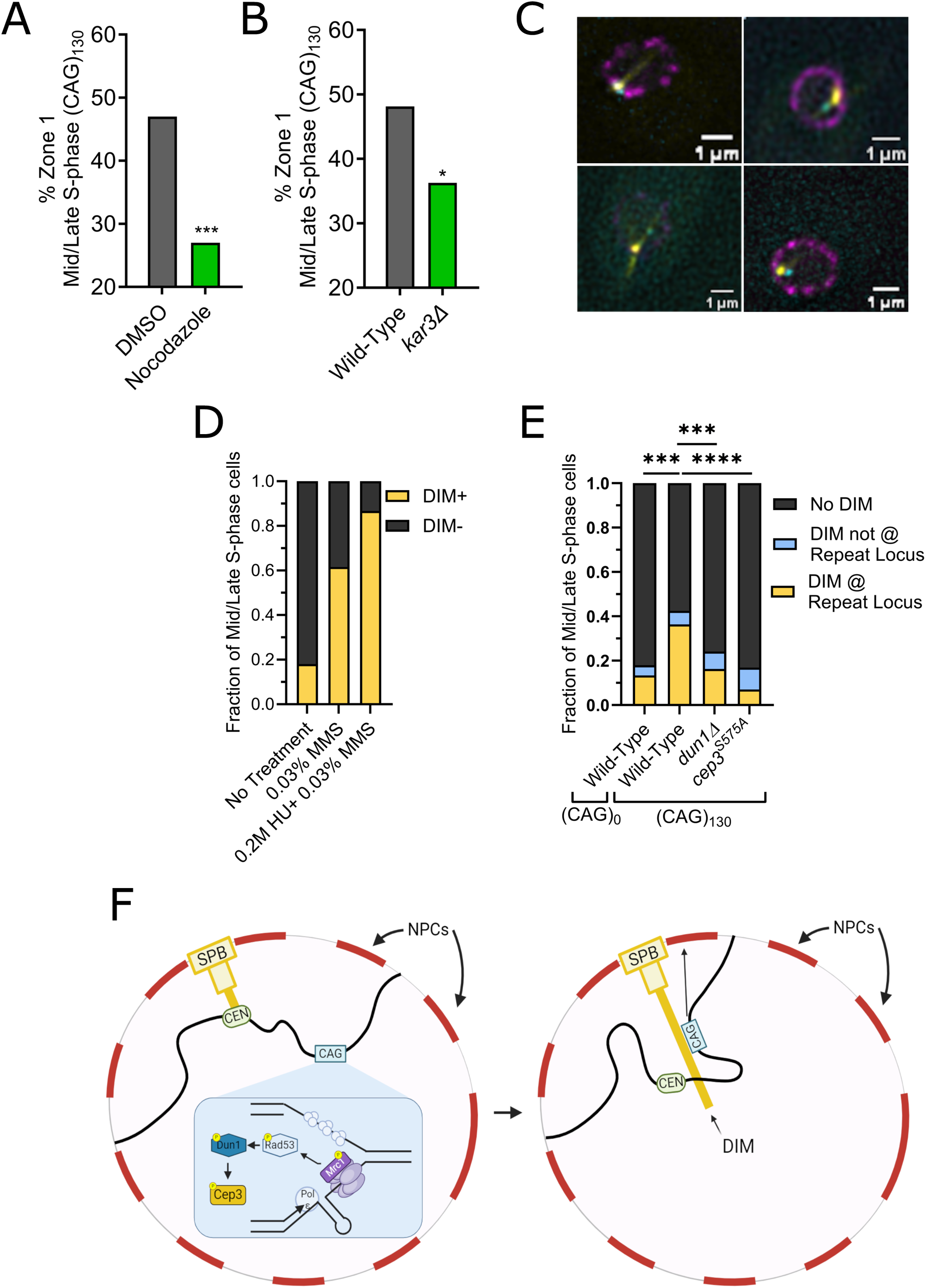
Damage-Inducible Microtubules associate with expanded CAG repeats in a Dun1 and Cep3 phosphorylation-dependent manner. A) Percentage of zone 1 (CAG)_130_ cells in mid/late S-phase. Cells were treated with either 15µg/mL nocodazole or an equivalent volume of DMSO (0.1%). B) Percent zone 1 (CAG)_130_ cells for cells lacking Kar3. C) Example images of CFP-LacI colocalizing with DIMs. Bottom left image is a cell without a DIM for comparison. The yellow channel was contrast-adjusted for clarity, raw images can be found in Figure S2A. D) Fraction of mid-late S phase cells (no CAG tract strain) treated as indicated with at least 1 DIM. E) Fraction of DIM containing cells of the indicated genotype. DIMs, if present, were scored as colocalized or not colocalized with the CFP-LacI focus. (*) P≤0.05, (***) P≤0.001, (****) P ≤ 0.0001 compared with wild-type by Fisher’s exact test (zoning assay) or chi-square test (DIM assay). See Table S1 (for zoning) or Table S4 (for DIM) for the exact number of cells analyzed, percentages, and P-values. F) Proposed model for the relocation of a collapsed fork at the CAG repeat to the nuclear pore complex. Collapsed or uncoupled forks at a CAG repeat result in a Mrc1 phosphorylation followed by Rad53 activation. Rad53 activated Dun1 leads to phosphorylation of Cep3. This phosphorylation alters the SPB-CEN attachment, leading to the formation of DIMs. This may allow repositioning of the centromere on the DIM, for example in a side-on attachment to permit relocation of the repeat to the NPC. Created with BioRender.com

To determine if DIMs were being recruited to the CAG repeat locus, we integrated an additional copy of the tubulin subunit Tub1 tagged with the YFP variant mVenus at the *TUB1* genomic locus ^58^. GFP-LacI was replaced with CFP-LacI and GFP-Nup49 with Nup49-mCherry to allow simultaneous visualization of all 3 nuclear components (Figure 4C). Consistent with previous data ^25^, treatment with 0.03% MMS caused DIM formation in 62% of S-phase cells (Figure 4D). To test whether DIMs were also induced by fork collapse, they were measured in cells treated with 0.03%MMS + 0.2M Hydroxyurea for 60 minutes, which causes genome-wide fork collapse and relocation of marked chromosome loci to the nuclear periphery ^1,11^. Under fork collapsing conditions, 87% of mid-to-late S phase cells contained a DIM (Figure 4D). We then tested whether a single expanded (CAG)_130_ repeat locus was sufficient to cause DIM formation in cells. We observed that there is an increase in prevalence of DIM formation in cells with an expanded CAG repeat to 36% compared to 18% for the no-tract control (Figure 4E). The majority of these DIMs (74%) colocalized with the repeat locus (Figure 4E; see Figures 4C and S2 examples). DIM formation was dependent upon Dun1 and Cep3 phosphorylation as deletion of Dun1 or mutation to *cep3^S575A^* caused a significant reduction of DIMs (Figure 4E). The repeat tract-induced DIMs were specifically affected in these mutants and were reduced to 7% in the *cep3^S575A^*strain (Figure 4E). We conclude that there is a checkpoint-dependent induction of DIMs and recruitment of a DIM to the site of fork collapse. These DIMs appear to be required to mediate relocation of the collapsed fork to the NPC as deletion of *KAR3* or treatment with nocodazole, both of which perturb DIM formation, led to a decrease in the frequency of relocation of the CAG tract to the nuclear periphery.

## Discussion

In this study we aimed to determine if the DNA damage checkpoint response plays a role in the relocation of collapsed forks to the nuclear pore complex (NPC). We previously showed that long CAG/CTG repeat tracts activate the DNA damage response and that the fork protection complex containing Mrc1 is crucial for preventing CAG fragility and instability and promoting fork progression through the repeat ^14,15,17^. Other systems showed varying checkpoint requirements for relocation of chromosomes with persistent DSBs, aphidicolin stalled forks or eroded telomeres ^1,3,22,23^. It was unknown what, if any, role the checkpoint played in the relocation of stalled/collapsed forks caused by expanded CAG repeats to the nuclear periphery. Even for types of damage where a requirement for checkpoint proteins had been established, the mechanism by which the checkpoint pathway mediates repositioning of damaged chromosomal loci had not been elucidated.

Our results indicate that the Mrc1/Rad53 axis is required for the repositioning of CAG repeats to the NPC in mid-to-late S phase. The (CAG)_130_ repeat forms hairpin structures that interfere with DNA replication ^59,60^ and CAG repeats have similar relocation requirements as cells which have been treated with both HU and MMS which causes fork collapse ^1,2,11^, thus it was speculated that the relocation event was a consequence of a collapsed fork ^2,11^. Since the phosphorylation of Mrc1 is required for the efficient sensing and stabilizing of stalled or uncoupled forks and the checkpoint activity of Mrc1 is critical for the relocation of the expanded repeat to the nuclear periphery, this implies that a stalled or uncoupled fork is the initial lesion being sensed and then processed for relocation to the NPC.

We found that there is redundancy in the sensing of the lesion caused by the (CAG)_130_ tract, suggesting that by-products of fork uncoupling such as ssDNA or an exposed DSB end caused by fork reversal or breakage also come into play. It was revealing that both the Mec1 and Rad9 pathways needed to be inactivated in order to abolish relocation to the NP. Long-range resection and RPA SUMOylation are required for CAG repeat relocation to the NPC ^11^, thus we would expect RPA-coated ssDNA, which recruits Mec1, to be available prior to relocation. Rad9, which is activated by both Mec1 and Tel1, was required only when Mec1 and/or the 9-1-1 complex are compromised, indirectly suggesting a role for Tel1. Tel1 has been shown to bind to the MRX complex ^61^, which binds DSB ends and is required for relocation ^11^. These results imply that either ssDNA or a DSB end can activate the pathway needed for repositioning, suggesting that there could be a transition from the initial uncoupled fork to a reversed or broken fork that provides a DSB end (Figure 1E). This redundancy in signaling leads to a robust mechanism to activate the Rad53 checkpoint in response to the initial fork stall caused by an expanded CAG repeat.

Rad53 is the master regulator of the checkpoint in *S. cerevisiae* and has multiple roles in the response to stalled or collapsed replication forks ^26,62–64^. One well-studied target of Rad53 is the kinase Dun1 ^37^, which we determined was needed for relocation (Figure 2B). It is important to note that while Dun1 is a required target of Rad53 for relocation, it may not be the only target as the kinase-deficient *rad53^K227A^* mutant is proficient for Dun1 activation ^65^, yet does not relocate. Dun1 phosphorylates several downstream targets including the recombination factor Rad55 ^37^, RNR inhibition factors (Crt1, Sml1, and Dif1 reviewed in ^38^), and the kinetochore protein Cep3 ^45^. Of these targets, the phosphorylation of Cep3 was required for relocation (Figure 2C).

The yeast centromere is only ∼120bp in length and has three sub-sections: CDEI, CDEII, and CDEIII. Kinetochore proteins at each centromere bind to a single microtubule, which is attached to the spindle pole body (SPB) throughout most of the cell cycle ^66^, with a transient release during centromere replication ^67^. Cep3 has a zinc finger DNA binding domain that interacts with the CDEIII region as a member of the CBF3 complex (reviewed in ^68^). Phosphorylation of the kinetochore has been widely studied and has varying functions (reviewed in ^69,70^. For example, phosphorylation of the kinetochore proteins, such as Dam1, by Aurora B kinase (Ipl1) destabilizes erroneous microtubule attachments in mitosis ^71,72^. However, the consequences of Cep3 phosphorylation have been comparatively poorly described. It was previously suggested that Cep3 phosphorylation led to a release of centromere constraint to allow the increased global mobility of chromosomes observed in response to an HO-induced DSB at the *MAT* locus in the G2/M phase of the cell cycle ^45^. However, another group was unable to show a Cep3-dependent increased radius of constraint ^50^ using similar conditions. Our results indicate that Cep3 phosphorylation may act through another mechanism besides increasing local or global chromatin mobility, at least in response to replication fork collapse. We propose that the DNA damage dependent phosphorylation of Cep3 is causing centromere release from the SPB attachment normally present during S phase. Consistent with this model, the requirement for Cep3 phosphorylation can be obviated by centromere inactivation (Figure 3A).

DIMs form in response to DNA damage induced by MMS, zeocin, camptothecin, or I-SceI-induced DSBs ^25^. With this work, we’ve now shown that while DIM formation is common after treatment with MMS, which stalls forks, it is dramatically increased after treatment with HU + MMS, which has been shown to be fork collapsing. DIM formation even occurs as a result of a single fork collapse caused by an expanded CAG repeat tract. Interestingly, much like how CEN inactivation eliminates the need for Cep3 phosphorylation for relocation, CEN inactivation can cause spontaneous formation of DIMs ^25^. This suggests that while the initial damage (whether a DSB or uncoupled/reversed/broken fork) is responsible for centromere release from the SPB microtubule through Cep3 phosphorylation, the DIM may form as a result of that release rather than directly responding to the damage. Thus, we hypothesize that release of the centromere from the end of the SPB microtubule triggers the formation of a DIM.

The microtubule motor protein Kar3 has been shown to promote the capture and directional movement of damaged loci by DIMs ^25^. When centromeres are inactivated, kinetochores that were previously attached to the microtubule plus end reposition and bind the microtubule side-on ^73,74^. Kar3, as a minus end directed motor, is involved in repositioning the centromere on the microtubule, by hindering tethering of the kinetochore to the plus end of the microtubule ^73–75^. During this kinetochore release and repositioning process, the damaged repeat locus could be freed up to move to the NPC. If this is the case, then we might expect that the presence of Kar3 could increase the duration of this “relocation permissive” state by antagonizing reattachment of the kinetochore to the microtubule plus end, which could explain why a *kar3Δ* reduces but does not completely abrogate relocation to the NPC. What remains unclear is whether the microtubule interacts directly with the collapsed fork, or if the DIM allows for repositioning of the lesion in an indirect manner, for example by allowing either free movement of the damaged locus or for the centromere to slide along the DIM. The former is an attractive hypothesis due to the increased colocalization of the repeat locus with the DIM (Figure 4E). It has recently been shown that DIMs serve as sites of repair center nucleation, with liquid droplets of Rad52 colocalizing with DIMs ^76^, and we’ve previously shown that many repair proteins, including Rad52 ^2^ and RPA ^11^ are recruited to the site of the collapsed fork prior to relocation. These factors have been shown to become SUMOylated, and their SUMOylation has been shown to be critically required for relocation to the NPC through an interaction with Slx5/8 ^11^. Slx5/8 in turn is critical for anchoring of damaged DNA at the NPC (reviewed in ^8^). SUMOylation has also been shown to promote condensation of repair factors ^77^. Possibly, SUMOylation or factors within SUMOylation-induced repair centers could play a role in the association between the collapsed fork at the CAG tract and the DIM.

Thus, we propose that after fork collapse at the CAG repeat, the DNA replication checkpoint results in the phosphorylation of Cep3. This phosphorylation causes centromere release from the SPB, which promotes DIM formation. The DIM allows movement of the collapsed fork locus to the periphery in a manner yet to be fully elucidated, but which may involve interaction of the damaged locus with the DIM. Once at the periphery, interactions between SUMOylated repair factors, Slx5/8, and the nucleoporins tether the locus to the NPC for repair or fork restart (Figure 4F).

There is prior evidence in other types of eukaryotic cells for a physical mechanism by which movement of lesions to repair centers within the nucleus occurs. Heterochromatic DSBs in *Drosophila* require nuclear F-actin and myosin for directed movement to the nuclear periphery ^78^. In human U2OS cells the relocation of collapsed forks caused by aphidicolin is dependent on the polymerization of nuclear F-actin, and late-replication foci were observed to move along actin filaments to the nuclear periphery ^3^. Further it’s been shown that recruitment of nuclear actin in response to damage can control pathway choice in response to replication stress ^79^. It is unclear if yeast have equivalent nuclear actin filaments or if directed movement to the NP would be detectable in the smaller yeast nucleus. However, in yeast, the kinesin-14 motor protein complex is required for movement of telomeric DSBs from their usual attachment site on the nuclear membrane to NPCs and these were observed to move along DIMs ^25,80^. Our results suggest that collapsed forks may also utilize DIMs to facilitate their movement to the yeast NPC. In human cells, it was recently reported that cytoplasmic microtubules that form in response to DSBs push on the nuclear envelope to form nuclear envelope tubules (NETs) that elongate into the nucleus and bring the nuclear envelope into the proximity of DSBs within the nuclear interior, and the DSBs may also move toward the dsbNETs ^81^. The formation of these dsbNETs serves to mobilize repair proteins to sites of DSBs and is dependent on the ATM/ATR/DNA-PK checkpoint response ^81^. Thus, the involvement of microtubules in mobilizing a repair response seems to be conserved, though the details of where in the nucleus the damage-inducible microtubules form and how they interact with the damaged locus may differ between organisms. It will be interesting to elucidate if DIMs are utilized in multicellular eukaryotic cells or if their function has been replaced by a combination of nuclear actin filaments and dsbNETs in these larger nuclei.

In summary, our results illuminate a new role for the DNA replication and DNA damage checkpoint response, which is to phosphorylate a kinetochore component to allow for repositioning of a collapsed fork within the nucleus. Since this repositioning is important for repair or fork restart, we identify it as a crucial new role for the DNA damage checkpoint response in *S. cerevisiae*. Repositioning of damaged loci to a repair center is present across many organisms. Where it has been studied, the checkpoint is critical to facilitate this relocation, but the targets involved have not been elucidated. Given the importance of recovery from replication fork collapse, a mechanism to allow centromere release to facilitate damaged locus repositioning might be especially critical at regions with fork-stalling DNA structures and for cancerous or aging cells experiencing high levels of replication stress.

## Acknowledgements

We would like to thank Stéphane Marcand for the gift of the *pGAL1*-CEN6 plasmid and Wei-Lih Lee for the *p*HIS3 *p*:Venus-Tub1+3’UTR::LEU2 plasmid.

## Author Contributions

JW: Conceptualization, Investigation, Analysis, Supervision, Writing

TM: Conceptualization, Investigation, Analysis, Software, Writing

MJ: Investigation, Analysis

CHF: Supervision, Resources, Writing, Administration, Conception

---

**Figure S1:**
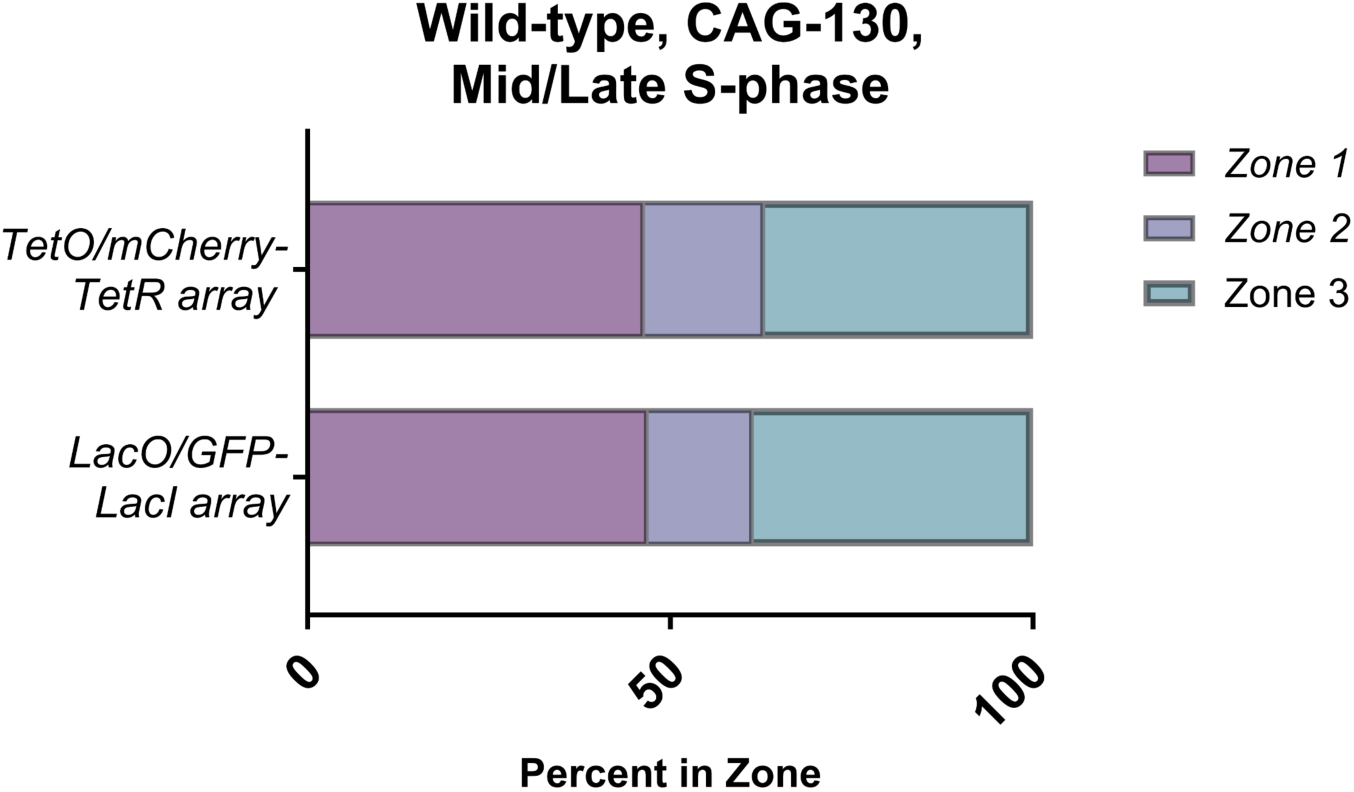
Comparison between wild-type strains. Percent of zone 1 foci for (CAG)_130_ mid/late S-phase cells for wild-type cells with the LacO/GFP-LacI array or TetO/mCherry-TetR array ∼6.4kb away from (CAG)_130_ near ARS0607 on Chr VI. The number of cells analyzed per strain were 157 and 162 respectively.

**Figure S2:**
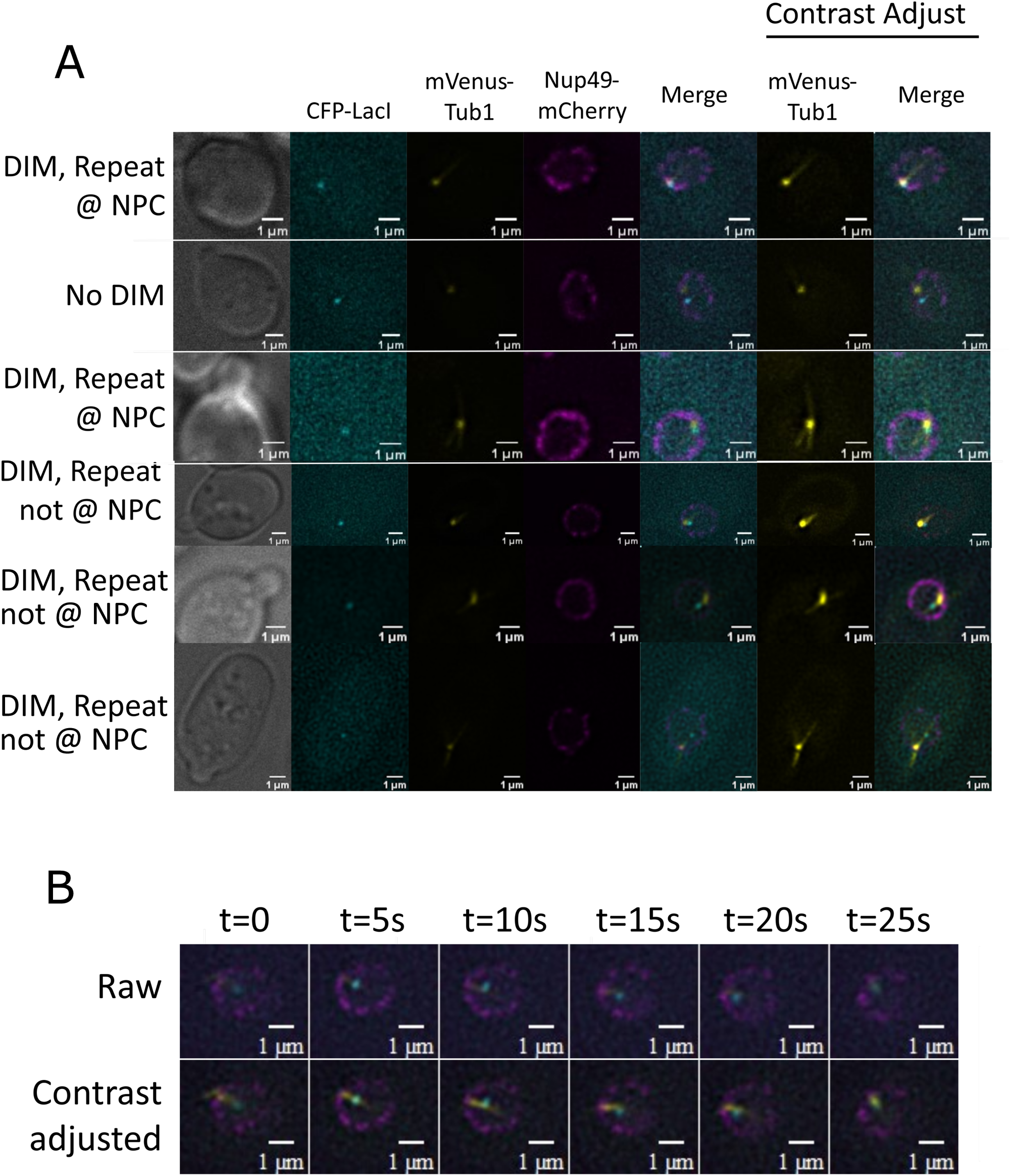
DIM Examples: A) Example images of Damage-inducible microtubules. Defined as monopolar microtubule extrusions during S-phase which exhibit rapidly decaying fluorescence intensity. Cell lacking a DIM shown for context. B) Example movie of a DIM associating with an expanded CAG repeat locus. Video acquired at 0.2 frame/s.

## Method Details

### Yeast strains and genetic manipulation

Standard procedures were used in cell growth and medium preparation. All *S. cerevisiae* strains were made in the W303 (RAD5+) background and are listed in Table S5. Point mutant alleles have been previously characterized: *rad53^K227A^*^82^, *cep3^S575A^*^45^. The point mutants were generated using CRISPR-Cas9 methods as described previously^83^. See Table S6 for primers used to create each mutation. Correct mutation was screened by both restriction enzyme digest and sequencing. The *P_GAL1_-CEN6* construct has been previously characterized ^51^. Centromere inactivation was confirmed by plating cells onto galactose containing media and observing a decrease in viability compared to glucose. Venus-Tub1 was introduced as an additional copy using plasmid pHIS3p:Venus-Tub1+3’UTR::LEU2, which was a gift from Wei-Lih Lee ^58^.

### Microscopy

Colonies were checked for 130-CAG repeats by PCR with primers flanking the repeat (CTGRev2/T720 or 2StepCAG_pAG32-F/R). PCR products separated on a 2% agarose gel (MetaPhor, Lonza). Cells from colonies with the correct repeat length were inoculated into YC media, grown overnight at 30C with shaking, diluted back to OD=0.2 and grown to approximately 5 × 10^6^ cells per mL in YC media. Cells were fixed with 4% paraformaldehyde and washed 3 times with 1X PBS. 1.4% YC agar was used to make agar pads on depression slides and 5 µL of cells were added on top with a coverslip. Z-stack images were taken using a Zeiss AX10 fluorescent microscope and/or a Delta Vision Ultra High-Resolution using an Olympus 100X with 1.45NA. Step interval size was 0.15-0.2um and approximately 25 Z-planes were taken per field of cells. Exposure time was approximately DIC: 100ms; GFP: 300ms and mCherry: 500ms (for strains with mCherry-TetR). Images were deconvolved, and three-zoning criteria was used to evaluate the location of the GFP foci for mid/late S-phase cells with the ImageJ point picker program as described in ^84^. Only the middle two-thirds of the stacks were used for analysis. Mid/Late S-phase cells were determined by yeast morphology using bud size criteria of 2/3^rd^ or less, and at least 15% the size of the mother cell ^85^. For drug treatments, cells were grown to approximately 1 × 10^7^, split into 2 cultures and grown for 30 minutes in either 0.1% dimethyl sulfoxide (DMSO) or 15ug/mL nocodazole. The cells were then fixed and imaged as described above. For DIM experiments, cells were prepared as described above and imaged with the filters (and exposure): Cyan (200ms), Yellow (200ms), Red (200ms). Cells were scored for colocalization determined by any overlap (not inclusive of “touching”) of the repeat and the nuclear periphery, and of the repeat with DIMs. DIMs were defined as monopolar microtubule structures emanating from the SPB during mid-late S phase (See Figure 4C for examples).

### *P_GAL1_*-CEN6 Inactivation

Colonies were checked for 130-CAG repeats by PCR with primers flanking the repeat (CTGRev2/T720). Cells from colonies with the correct repeat length were grown overnight in synthetic complete media +2% glucose. Cultures were spun down and resuspended in YEP-lactate media to an OD_600_ of 0.3 and grown overnight. Cultures were spun down, split in half, and resuspended in synthetic complete media +2% galactose or +2% glucose to an OD_600_ of 0.3. Cultures were grown for 3 hours followed by the zoning assay protocol as outlined above.

### MSD Analysis

Colonies were checked for 130-CAG repeats by PCR with primers flanking the repeat (2StepCAG_pAG32-F/R). Cells with correct repeat length were grown to approximately OD_600_ of 0.5 before being transferred to a concanavalin A-coated chamber slide for attachment. From there, cells were synchronized by addition of alpha factor (1µM final concentration) then released by washing 3X with YPD media. A single field of cells was imaged with DeltaVision Ultra High Resolution microscope using a 100X objective using optical axis integration every 1.5s for 5 minutes. This was repeated using cells in different areas of the slide until a total of 100 minutes of timepoints were taken. Exposure was: Red 400ms, Green 400ms, Pol 50ms (every 50^th^ frame). Images were analyzed with CellProfiler, tracking the LacO/LacI-mCherry focus and correcting for nuclear translation by tracking Nup49-GFP. Tracked foci were quantified using 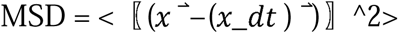 and radius of constraint = sqrt(5/4 x MSD plateau) (Dion & Gasser, 2013). The plateau was calculated by regression in Graphpad PRISM software using the first 100 Δt timepoints.

### Quantification and statistical analysis

Prism software was used to calculate statistical significance. For microscopy experiments Fisher’s exact test (Relocation experiments) or chi square test (DIM experiments) were used.

### Data and software availability

All raw data is available in the supplemental tables

