## Supplemental Tables for "The DNA Replication Checkpoint Targets the Kinetochore for Relocation of Collapsed Forks to the Nuclear Periphery"

**Table 1: Zoning assay data**

| **Strain** | **Strain No.** | **Zone 1 foci per strain** | | | **Total Zone 1 foci** | | | **p-value (Fisher's exact to WT unless otherwise indicated)** |
| --- | --- | --- | --- | --- | --- | --- | --- | --- |
|  |  | **No.** | **%** | **Total No. cells per strain** | **No.** | **%** | **Total No. cells** |  |
| Wild-type CAG-130 | 2744 | 78 | 48.1 | 162 | 78 | 48.1 | 162 | ----- |
| *slx5Δ* | 2797 | 46 | 30 | 152 | 46 | 30 | 152 | 0.001 |
| *chk1Δ* | 5133/5134 | 61 | 40.7 | 150 | 61 | 40.7 | 150 | 0.21 |
| *rad53Δsml1Δ* | 5203 | 29 | 30.5 | 95 | 45 | 30 | 150 | 0.001 |
|  | 5219 | 16 | 29 | 55 |  |  |  |  |
| *rad53^K227A^* | 4952 | 24 | 27.3 | 88 | 49 | 30 | 163 | 0.001 |
|  | 4953 | 25 | 33.3 | 75 |  |  |  |  |
| *mec1Δsml1Δ* | 3447 | 73 | 48 | 152 | 73 | 48 | 152 | 1 |
| *ddc1Δ* | 4214 | 56 | 43.8 | 128 | 70 | 45.3 | 155 | 0.65 |
|  | 4215 | 14 | 51.9 | 27 |  |  |  |  |
| *rad24Δ* | 4120 | 22 | 45.8 | 48 | 53 | 43.4 | 122 | 0.47 |
|  | 4121 | 31 | 41.8 | 74 |  |  |  |  |
| *rad9Δ* | 3762 | 58 | 47.5 | 122 | 74 | 45.7 | 169 | 0.44 |
|  | 3763 | 16 | 40 | 40 |  |  |  |  |
| *tel1Δ* | 3582 | 34 | 45.3 | 75 | 68 | 45.3 | 150 | 0.65 |
|  | 3583 | 34 | 45.3 | 75 |  |  |  |  |
| *mec1Δrad9Δ sml1Δ* | 4913 | 27 | 36.5 | 74 | 54 | 35.1 | 154 | 0.02 |
|  | 4926 | 27 | 33.8 | 80 |  |  |  |  |
| *rad9Δrad24Δ* | 4096 | 39 | 32.5 | 120 | 50 | 33.1 | 151 | 0.008 |
|  | 4097 | 11 | 35.5 | 31 |  |  |  |  |
| *mec1Δrad9Δ rad24Δsml1Δ* | 5093 | 17 | 21.3 | 80 | 39 | 24.4 | 160 | 0.0001 |
|  | 5094 | 22 | 27.5 | 80 |  |  |  |  |
| *rad9Δtel1Δ* | 5193 | 34 | 45.3 | 75 | 72 | 47.4 | 152 | 0.91 |
|  | 5194 | 38 | 49.4 | 77 |  |  |  |  |
| *rad9Δrad24Δ tel1Δ* | 5211 | 22 | 28.9 | 76 | 49 | 32.5 | 151 | 0.006 |
|  | 5212 | 27 | 35.5 | 75 |  |  |  |  |
| *dun1Δ* | 5135/5136 | 38 | 27.3 | 139 | 38 | 27.3 | 139 | 0.0002 |
| *rad55Δ* | 5198 | 26 | 43.3 | 60 | 77 | 46.7 | 165 | 0.83 |
|  | 5199 | 44 | 44 | 50 |  |  |  |  |
|  | 5200 | 29 | 52.7 | 55 |  |  |  |  |
| *rad55Δrad57Δ* | 5226 | 40 | 54.1 | 74 | 82 | 53.9 | 152 | 0.31 |
|  | 5227 | 42 | 53.8 | 78 |  |  |  |  |
| *sml1Δ* | 5201 | 26 | 50.9 | 51 | 73 | 47.4 | 154 | 0.91 |
|  | 5202 | 47 | 45.6 | 103 |  |  |  |  |
| *dun1Δsml1Δ* | 5305 | 11 | 22.4 | 49 | 45 | 29.6 | 152 | 0.0008 |
|  | 5306 | 34 | 33 | 103 |  |  |  |  |
| *cep3^S575A^* | 5277 | 25 | 32.9 | 76 | 58 | 33.3 | 174 | 0.008 |
|  | 5287 | 33 | 33.7 | 98 |  |  |  |  |
| *cep3^S575E^* | 5393 | 60 | 58.2 | 103 | 95 | 57.9 | 164 | 0.09 |
|  | 5395 | 35 | 57.4 | 61 |  |  |  |  |
| *dun1Δcep3^S575E^* | 5379 | 53 | 51.5 | 103 | 85 | 55 | 154 | 0.22 |
|  | 5380 | 32 | 62.7 | 51 |  |  |  |  |
| *cep3-S575A pGAL1-CEN6 -Galactose* | 5402 | 39 | 31.2 | 125 | 58 | 30.1 | 193 | 0.0007 |
|  | 5403 | 19 | 27.9 | 68 |  |  |  |  |
| *cep3-S575A pGAL1-CEN6 +Galactose* | 5402 | 51 | 42.5 | 120 | 86 | 48 | 179 | 1 |
|  | 5403 | 35 | 59.3 | 59 |  |  |  | 0.0004 compared to -Galactose |
| *nhp6aΔ* | 5297 | 34 | 45.3 | 75 | 87 | 45.1 | 193 | 0.59 |
|  | 5298 | 53 | 44.9 | 118 |  |  |  |  |
| *nhp6bΔ* | 5318 | 43 | 54.4 | 79 | 78 | 50.6 | 154 | 0.74 |
|  | 5319 | 35 | 46.7 | 75 |  |  |  |  |
| *nhp6aΔnhp6bΔ* | 5300 | 43 | 46.2 | 93 | 68 | 43.6 | 156 | 0.43 |
|  | 5301 | 25 | 39.7 | 63 |  |  |  |  |
| *mrc1Δ+pMRC1-AQ* | 5618 | 46 | 28.6 | 161 | 93 | 28.1 | 331 | 1.4E-5 |
|  | 5619 | 57 | 27.7 | 170 |  |  |  |  |
| *mad2Δ* | 5617 | 73 | 52.1 | 140 | 73 | 52.1 | 140 | .56 |
| *kar3Δ* | 5757 | 35 | 38.0 | 92 | 70 | 36.2 | 193 | 0.03 |
|  | 5758 | 35 | 34.7 | 101 |  |  |  |  |
| Wild Type CAG-130 + DMSO | 3116 | 76 | 46.9 | 162 | 76 | 46.9 | 162 | .91 compared to untreated |
| Wild Type CAG-130 +  15ug/mL Nocodazole | 3116 | 39 | 27.1 | 144 | 39 | 27.1 | 144 | .0002 compared to DMSO treated |

**Table 2: Zoning assay time course analysis of *cep3^S575E^***

|  | | | **Strain** | |
| --- | --- | --- | --- | --- |
|  |  |  | **Wild-type** | **cep3^S575E^** |
| **Time after alpha-factor release** | **20** | **No. Zone 1 foci** | 44 | 51 |
|  |  | **%** | 21.8 | 33.6 |
|  |  | **Total No. cells** | 202 | 152 |
|  |  | **p-value to WT*** | ---- | 0.02 |
|  | **30** | **No. Zone 1 foci** | 57 | 52 |
|  |  | **%** | 32 | 34.7 |
|  |  | **Total No. cells** | 179 | 150 |
|  |  | **p-value to WT*** | ---- | 0.63 |
|  | **40** | **No. Zone 1 foci** | 47 | 59 |
|  |  | **%** | 26.9 | 45.4 |
|  |  | **Total No. cells** | 175 | 130 |
|  |  | **p-value to WT*** | ---- | 0.001 |
|  | **50** | **No. Zone 1 foci** | 71 | 93 |
|  |  | **%** | 34 | 52.8 |
|  |  | **Total No. cells** | 206 | 176 |
|  |  | **p-value to WT*** | ---- | 0.0004 |
|  | **60** | **No. Zone 1 foci** | 89 | 80 |
|  |  | **%** | 48.9 | 52.9 |
|  |  | **Total No. cells** | 182 | 151 |
|  |  | **p-value to WT*** | ---- | 0.51 |

*using the Fisher's exact test

**Table 3: Mean Square Displacement (MSD) analysis data**

|  | cep3-S575A | | | Wild-Type | | |
| --- | --- | --- | --- | --- | --- | --- |
| min post G1 release | Radius of constraint (μm)* | SEM | N | Radius of constraint (μm)* | SEM | N |
| 5 | 0.6599 | 0.07012 | 5 | 0.746492 | 0.06649 | 9 |
| 10 | 0.9039 | 0.1217 | 4 | 0.619274 | 0.1205 | 6 |
| 15 | 0.8986 | 0.1078 | 3 | 0.966566 | 0.06725 | 3 |
| 20 | 0.7989 | 0.2769 | 2 | 0.700625 | 0.07287 | 5 |
| 25 | 0.9523 | 0.1002 | 5 | 0.631862 | 0.07965 | 7 |
| 30 | 0.8444 | 0.1185 | 6 | 0.689656 | 0.09995 | 3 |
| 35 | 0.5941 | 0.08926 | 6 | 0.652495 | 0.08602 | 8 |
| 40 | 0.8201 | 0.07469 | 8 | 0.707284 | 0.04335 | 7 |
| 45 | 0.8821 | 0.06559 | 5 | 0.729812 | 0.07197 | 14 |
| 50 | 0.7688 | 0.1222 | 8 | 0.745822 | 0.04515 | 8 |
| 55 | 0.8986 | 0.07013 | 12 | 0.744228 | 0.06408 | 7 |
| 60 | 0.778 | 0.0777 | 8 | 0.735102 | 0.07607 | 5 |
| 65 | 0.6851 | 0.06772 | 3 | 0.732462 | 0.03682 | 5 |
| 70 | 1.045 | 0.1012 | 4 | 0.649808 | 0.1075 | 4 |
| 75 | 0.7842 | 0.08749 | 3 | 0.618971 | 0.06715 | 10 |
| 80 | 0.9161 | 0.1144 | 3 | 0.700179 | 0.06497 | 7 |
| 85 | 0.5469 | 0 | 1 | 0.677311 | 0.2221 | 2 |
| 90 | 0.6527 | 0.1263 | 3 | 0.809398 | 0.05053 | 3 |
| 95 | 0.6576 | 0.06694 | 6 | 0.6257 | 0.06277 | 4 |
| 100 | 0.8234 | 0.06687 | 4 | 0.693001 | 0.07622 | 4 |

*calculated from the first Δt =[0,100] seconds of the 5 minute interval shown; cells were imaged every 1.5 seconds.

**Table 4: Damage-Inducible Microtubule (DIM) data.**

| **Strain** | **Strain No.** | **No. cells with DIM @ Repeat** | **No. cells with DIM not @ Repeat** | **No. cells with no DIM** | **p value (compared to WT, CAG-130)** |
| --- | --- | --- | --- | --- | --- |
| WT, CAG-130 | 5779 | 40 | 14 | 95 | N/A |
| WT, No repeat | 5805 | 20 | 7 | 123 | p < 0.0001 |
| *dun1Δ* | 5806 | 11 | 5 | 47 | p = 0.0008 |
| *dun1Δ* | 5807 | 12 | 6 | 60 |  |
| *cep3-S575A* | 5920 | 8 | 10 | 90 | p < 0.0001 |
| *cep3-S575A* | 5921 | 5 | 8 | 63 |  |

| **WT Strain No.** | **Treatment** | **DIM** | **No DIM** | **p value (compared to no treatment) chi-square** |
| --- | --- | --- | --- | --- |
| 5805 | No Treatment | 27 | 123 | N/A |
| 5805 | 0.03% MMS | 98 | 61 | <0.0001 |
| 5805 | 0.03% MMS + 0.2M HU | 65 | 10 | <0.0001 |

**Table 5: Strains used in this study**

| **CFY Strain Number** | **Strain Background** | **Genotype** | **Reference** |
| --- | --- | --- | --- |
| 2744 | W303 | *MATa ade2-1 can1-100 his3-11,-15::GFP-LacI:HIS3 trp1-1, ura3-1, leu2-3,-112 nup49::GFP-NUP49, Chr6int1::lacO:4xLexA array:TRP1, Chr6int2::CAG130:HPH* (*Chr6int1* is 272 bp upstream of *ARS607,* within *PES4*) (Chr6int2 is 6371 bp from Chr6int1 and 560bp upstream of Ta(AGC)F gene) | Su et al. 2015 |
| 2797 | W303 | *CFY2744; slx5::KANMX* | Su et al. 2015 |
| 5133 | W303 | *CFY2744; chk1::KANMX* | This study |
| 5134 |  |  |  |
| 3116 | W303 | *MATa*, *RAD52-YFP, GFP-NUP49, ade2::TetR-mCherry-ADE2, lys5::LacI-CFP-TRP1, leu2::LoxP, ZWF1:cutsite* (Lmn::lys5::IsceIcs::LEU2::LacO array::Lmn), met10::lmn adaptamers::HIS3::TetOps-LexA.; Chr6int2::CAG130-HPH is 560 bp upstream of tA(AGC)F) | Su et al. 2015 |
| 5201 | W303 | *CFY3116; sml1::NATMX* | This study |
| 5202 |  |  |  |
| 5203 | W303 | *CFY5202; rad53::KANMX* | This study |
| 5219 |  |  |  |
| 4952 | W303 | *CFY2744; rad53^K227A^-KANMX* | This study |
| 4953 |  |  |  |
| 3426 | W303 | *CFY2744; sml1::KANMX* | Su et al. 2015 |
| 3446 | W303 | *CFY3426; mec1::KANMX* | Su et al. 2015 |
| 3447 |  |  |  |
| 4214 | W303 | *CFY2744; ddc1::KANMX* | This study |
| 4215 |  |  |  |
| 4120 | W303 | *CFY2744; rad24::NATMX* | This study |
| 4121 |  |  |  |
| 3762 | W303 | *CFY2744; rad9::KANMX* | This study |
| 3763 |  |  |  |
| 3582 | W303 | *CFY2744; tel1::KANMX* | Su et al. 2015 |
| 3583 |  |  |  |
| 2961 | W303 | *WT W303 Mat alpha* | Gasser Lab |
| 4525 | W303 | *CFY2744xCFY2961; Mat alpha, zoning assay cassette** | This study |
| 4889 | W303 | *CFY3582xCFY4525; tel1::KANMX, rad9::NATMX, zoning assay cassette** | This study |
| 4913 | W303 | *CFY4889xCFY3446; mec1::KANMX, rad9::NATMX, sml1::NATMX, zoning assay cassette** | This study |
| 4926 |  |  |  |
| 4096 | W303 | *CFY3762; rad24::NATMX* | This study |
| 4097 |  |  |  |
| 5018 | W303 | *CFY3762xCFY3446; mec1::KANMX, sml1::NATMX, rad9::KANMX, zoning assay cassette** | This study |
| 5093 | W303 | *CFY5018xCFY4096 mec1::KANMX, rad9::NATMX, rad24::LEU2, sml1::NATMX, zoning assay cassette** | This study |
| 5094 |  |  |  |
| 5193 | W303 | *CFY3582; rad9::NATMX* | This study |
| 5194 |  |  |  |
| 5211 | W303 | *CFY5193; rad24::LEU2* | This study |
| 5212 |  |  |  |
| 5135 | W303 | *CFY2744; dun1::KANMX* | This study |
| 5136 |  |  |  |
| 5198 | W303 | *CFY2744; rad55::KANMX* | This study |
| 5199 |  |  |  |
| 5200 |  |  |  |
| 5226 | W303 | *CFY5198; rad57::NATMX* | This study |
| 5227 |  |  |  |
| 5305 | W303 | *CFY5135; sml::NATMX* | This study |
| 5306 |  |  |  |
| 5277 | W303 | *CFY3116; cep3^S575A^* | This study |
| 5287 |  |  |  |
| 5393 | W303 | *CFY3116; cep3^S575E^* | This study |
| 5395 |  |  |  |
| 5379 | W303 | *dun1Δcep3^S575E^* | This study |
| 5380 |  |  |  |
| 5402 | W303 | *CFY5287; KANr-pGAL1-CEN6* | This study |
| 5403 |  |  |  |
| 5297 | W303 | *CFY2744; nhp6a::KANMX* | This study |
| 5298 |  |  |  |
| 5318 | W303 | *CFY2744; nhp6b::NATMX* | This study |
| 5319 |  |  |  |
| 5300 | W303 | *CFY5298; nhp6b::NATMX* | This study |
| 5301 |  |  |  |
| 5601 | W303 | *CFY3116; mrc1::KANMX* | This study |
| 5602 |  |  |  |
| 5616 | W303 | *CFY3116; mad2Δ* | This study |
| 5617 |  |  |  |
| 5757 | W303 | *CFY3116; kar3::KANMX* | This study |
| 5758 |  |  |  |
| 5618 | W303 | *CFY5601; pAO138-MRC1-AQ-URA3* | This study |
| 5619 | W303 | *CFY5602; pAO138-MRC1-AQ-URA3* | This study |
| 5779 | W303 | *MATalpha; ade2-1 can1-100 his3-11,-15, trp1-1, ura3-1::CFP-LacI:URA3, leu2-3,-112 nup49:: NUP49-mCherry:HpH, Chr6int1::lacO:4xLexA array:TRP1, Chr6int2::CAG130* (*Chr6int1* is 272 bp upstream of *ARS607,* within *PES4*) (Chr6int2 is 6371 bp from Chr6int1 and 560bp upstream of Ta(AGC)F gene) Tub1:3’UTR-mVenus-Tub1:LEU2 | This study |
| 5805 | W303 | *MATa; ade2-1 can1-100 his3-11,-15, trp1-1, ura3-1::CFP-LacI:URA3, leu2-3,-112 nup49:: NUP49-mCherry:HpH, Chr6int1::lacO:4xLexA array:TRP1,* (*Chr6int1* is 272 bp upstream of *ARS607,* within *PES4*) Tub1:3’UTR-mVenus-Tub1:LEU2 | This study |
| 5806 | W303 | *CFY5779; dun1::KANMX* | This study |
| 5807 | W303 |  |  |
| 5920 | W303 | *CFY5779; cep3-S575A* | This study |
| 5921 | W303 |  |  |

*Zoning assay cassette: his3-11,-15::GFP-LacI:HIS3 nup49::GFP-NUP49 Chr6int1::lacO:4xLexA array:TRP1 Chr6int2::CAG130:HPH (Chr6int1 is 272 bp upstream of ARS607, within PES4) (Chr6int2 is 6371 bp from Chr6int1 and 560bp upstream of Ta(AGC)F gene)

**Table 6:** **Primers used in this study**

| **Primer sequence** | **Source** | **Identifier** |
| --- | --- | --- |
| 5’-CCTCAGCCTGGCCGAAAGAAAGAAA-3’ | Eton Bioscience Inc. | NewCAGfor |
| 5’-CAGTCACGACGTTGTAAAACGACGG-3’ | Eton Bioscience Inc. | NewCAGrev |
| 5’-GGTCAGGGCCTCAGCCTGGCCGAAA-3’ | Eton Bioscience Inc. | 2Step_pAG32_F |
| 5’-GGCCGGCGGAACGGGGCTCGAA-3’ | Eton Bioscience Inc. | 2Step_pAG32_R |
| 5'-TAATACGACTCACTATAGGG-3’ | Eton Bioscience Inc. | T7-20 |
| 5'-CCCAGGCCTCCAGTTTGC-3’ | Eton Bioscience Inc. | CTG rev2 |
| 5'-AACTACTGGGAAAACATTCG gttttagagctagaaatagcaagttaaaataagg-3' | Eton Bioscience Inc. | Rad53_pRCC_F |
| 5'-CGAATGTTTTCCCAGTAGTT tccgatcatttatctttcactgcggag-3' | Eton Bioscience Inc. | Rad53_pRCC_R |
| 5'-TTGCCACAGTAAAGAAAGCCATTGAAA GAACTACTGGGAAAACATTCGCCGTGGCGATTATAAGTAAACGCAAAGTAATAGGCAATATGGATGGTGTGAC-3' | Thermoscientific | Rad53^K227A^ Repair Template_F |
| 5'-GTCACACCATCCATATTGCCTATTACTT TGCGTTTACTTATAATCGCCACGGCGAATGTTTTCCCAGTAGTTCTTTCAATGGCTTTCTTTACTGTGGCAA-3' | Thermoscientific | Rad53^K227A^ Repair Template_R |
| 5'-TAGACAAGAATCGTTGCTTG gttttagagctagaaatagcaagttaaaataagg-3' | Eton Bioscience Inc. | Cep3-pRCC-F |
| 5'-CAAGCAACGATTCTTGTCTA tccgatcatttatctttcactgcgg-3' | Eton Bioscience Inc. | Cep3-pRCC-R |
| 5'-CGGGTCTTTGGTACCGTTGAATAAGC TTAGACAAGAAGCGTTGCTTGAAGAAGAGGACGAAAACAATACGGAACCAAGTGACTTCAGAACTATTGTAGAA-3' | Thermoscientific | Cep3^S575A^ Repair Template_F |
| 5'-TTCTACAATAGTTCTGAAGTCACTTGGT TCCGTATTGTTTTCGTCCTCTTCTTCAAGCAACGCTTCTTGTCTAAGCTTATTCAACGGTACCAAAGACCCG-3' | Thermoscientific | Cep3^S575A^ Repair Template_R |
| 5'-CGGGTCTTTGGTACCGTTGAATAAGC TTAGACAAGAAGAGTTGCTTGAAGAAGAGGACGAAAACAATACGGAACCAAGTGACTTCAGAACTATTGTAGAA-3' | Thermoscientific | Cep3^S575E^ Repair Template_F |
| 5'-TTCTACAATAGTTCTGAAGTCACTTGGTT CCGTATTGTTTTCGTCCTCTTCTTCAAGCAACTCTTCTTGTCTAAGCTTATTCAACGGTACCAAAGACCCG-3' | Thermoscientific | Cep3^S575E^ Repair Template_R |
| 5’- GTATTGAAAACCACTTCAAAGGGGCCCAATAGCACATTTATAGAATTCAAGTCaaaaaaaaaaaaaaaGACGGAAGTCCA-3’ | Eton Bioscience Inc | mad2Δ Repair Template |
| 5’- GGGCAGAAAGGTAACGCTTG gttttagagctagaaatagcaagttaaaataagg-3’ | Eton Bioscience Inc | Mad2-pRCC-F |
| 5’-CAACGTTACCTTTCTGCCC tccgatcatttatctttcactgcgg-3’ | Eton Bioscience Inc | Mad2-pRCC-R |
